# WNT4 regulates cellular metabolism via intracellular activity at the mitochondria in breast and gynecologic cancers

**DOI:** 10.1101/2022.07.15.499747

**Authors:** Joseph L. Sottnik, Madeleine T. Shackleford, Fabian R. Villagomez, Shaymaa Bahnassy, Steffi Oesterreich, Junxiao Hu, Zeynep Madak-Erdogan, Rebecca B. Riggins, Bradley R. Corr, Linda S. Cook, Lindsey S. Treviño, Benjamin G. Bitler, Matthew J. Sikora

## Abstract

Wnt ligand WNT4 is critical in female reproductive tissue development, with WNT4 dysregulation linked to related pathologies including breast cancer (invasive lobular carcinoma, ILC) and gynecologic cancers. WNT4 signaling in these contexts is distinct from canonical Wnt signaling yet inadequately understood. We previously identified atypical intracellular activity of WNT4 (independent of Wnt secretion) regulating mitochondrial function, and herein examine intracellular functions of WNT4. We further examine how convergent mechanisms of *WNT4* dysregulation impact cancer metabolism. In ILC, *WNT4* is co-opted by estrogen receptor α (ER) via genomic binding in *WNT4* intron 1, while in gynecologic cancers, a common genetic polymorphism (rs3820282) at this ER binding site alters *WNT4* regulation. Using proximity biotinylation (BioID), we show canonical Wnt ligand WNT3A is trafficked for secretion, but WNT4 is localized to the cytosol and mitochondria. We identified DHRS2, mTOR, and STAT1 as putative WNT4 cytosolic/mitochondrial signaling partners. Whole metabolite profiling, and integrated transcriptomic data, support that WNT4 mediates metabolic reprogramming via fatty acid and amino acid metabolism. Further, ovarian cancer cell lines with rs3820282 variant genotype are WNT4-dependent and have active WNT4 metabolic signaling. In protein array analyses of a cohort of 103 human gynecologic tumors enriched for patient diversity, germline rs3820282 genotype is associated with metabolic remodeling. Variant genotype tumors show increased AMPK activation and downstream signaling, with the highest AMPK signaling activity in variant genotype tumors from non-White patients. Taken together, atypical intracellular WNT4 signaling, in part via genetic dysregulation, regulate the distinct metabolic phenotypes of ILC and gynecologic cancers.

**Significance:** WNT4 regulates breast and gynecologic cancer metabolism via a previously unappreciated intracellular signaling mechanism at the mitochondria, with WNT4 mediating metabolic remodeling. Understanding WNT4 dysregulation by estrogen and genetic polymorphism offers new opportunities for defining tumor biology, precision therapeutics, and personalized cancer risk assessment.

## INTRODUCTION

The Wnt ligand WNT4, considered a “problem child” among Wnts [1, 2], is broadly critical for organogenesis and development. Complete loss-of-function is lethal *in utero* due to multi-organ dysgenesis/agenesis [1].

Beyond fetal development, WNT4 has mechanistically distinct roles in female reproductive tissues, in organogenesis of the ovaries and uterus, female sex differentiation, and pregnancy phenotypes in the uterus and mammary gland. Accordingly, WNT4 dysfunction is associated with reproductive, gynecologic, and endocrine pathologies including pre-cancerous uterine lesions, ovarian cancer, and the breast cancer subtype invasive lobular carcinoma (ILC) (reviewed in [1]). Though WNT4 engages cell- and tissue-specific downstream pathways that are not well defined, converging phenotypes in reproductive tissues and associated cancers suggest that understanding WNT4 regulation and signaling can provide new insights into cancers primarily affecting women.

In the mammary gland, WNT4 is critical for ductal elongation and branching during pregnancy. In this context, *WNT4* expression is induced by progesterone in progesterone receptor (PR)-positive luminal cells, and WNT4 protein subsequently signals in a paracrine manner, mediating stem cell proliferation and renewal likely via β- catenin-dependent signaling [3]. We found that in ILC cell lines, *WNT4* expression is co-opted and directly controlled by estrogen receptor α (ER), and that WNT4 is necessary for ER-driven growth and anti-estrogen resistance in ILC cell lines [4–6]. However, owning to hallmark loss of E-cadherin [7], ILC cells and tumors typically lack β-catenin protein and fail to engage β-catenin-dependent Wnt signaling [5, 7]. We found that ER-driven WNT4 instead controls mTOR signaling and mitochondrial dynamics in ILC cells; WNT4 knockdown compromises respiratory capacity leading to cell death [6]. This phenotype is putatively linked to the unique metabolism of ILC, which are considered metabolically quiescent vs invasive ductal carcinoma (IDC, i.e. breast cancer of no special type). Clinical ILC imaging shows reduced glucose uptake, as ILC tumors have ∼40% reduced glucose uptake vs IDC in ^18^F-fluorodeoxyglucose (FDG)/PET-CT imaging [8–13], and ILC tumors with no detectable FDG uptake are common especially in metastatic ILC [9, 13, 14]. Compared to IDC, expression of glucose metabolism genes are reduced in ILC [15], and tissue microarray studies show ILC express lower levels of related proteins (e.g. GLUT1/*SLC2A1*) [16]. Conversely, ILC tumors show differential expression of lipid metabolism-related proteins [15, 17], and accordingly, studies with anti-estrogen resistant ILC models identify increased lipid and glutamate metabolism as critical to the resistant phenotype [18–20]. Taken together, these observations suggest that ER+ ILC cells and tumors have a distinct metabolic phenotype compared to other breast cancers, which our work links directly to WNT4-dependent signaling activity. With cancer metabolism increasingly tractable as a treatment target, there is an urgent need to better define WNT4-mediated mechanisms of metabolic reprogramming.

Paralleling the essential role of WNT4 in mammary gland development, WNT4 dysregulation is associated with female-to-male sex reversal phenotypes, uterine agenesis, and ovarian dysgenesis [1]. Strikingly, over 20 genome-wide association studies (GWAS) link single nucleotide polymorphisms (SNPs) at the *WNT4* locus to increased risk of gynecologic pathologies including endometriosis and ovarian cancer [1]. Fine mapping and functional studies identify SNP rs3820282, a C>T transition in *WNT4* intron 1, as the likely pathogenic SNP [21, 22]. Of note, the rs3820282 variant allele frequency (VAF) varies from 0% to over 50% across racial/ethnic populations ([23], see Discussion). The rs3820282 variant creates a binding site for nuclear receptor-class transcription factors, also converting a half estrogen-response-element (ERE) to a full consensus ERE [5, 22]. Importantly, knock-in of the variant allele in mice is sufficient to increase *Wnt4* expression in gynecologic tissue *in vivo* [24]. We also showed that this locus is directly bound by ER in ILC cells (wild-type for rs3820282) but not other breast cancer cells [5]. These observations suggest convergent mechanisms of dysregulated *WNT4* gene expression, via ER and genetic polymorphism, respectively drive breast and gynecologic pathologies.

Toward understanding WNT4 signaling, we previously showed in ILC and ovarian cancer cells that WNT4 signals independent of canonical Wnt secretion and paracrine activity via an atypical intracellular mechanism [6, 25]. In this study, we used proximity biotinylation to profile WNT4’s interactome, signaling partners, and intracellular localization. We also used global untargeted mass spectrometry-based metabolomics analyses to profile the metabolic effects of WNT4 regulation and signaling. We performed reverse phase protein array (RPPA) analyses of 103 primary human gynecologic tumors, enriched for patient diversity and germline rs3820282 variant genotype, to determine the impact of the rs3820282/*WNT4* axis on tumor biology. These analyses corroborate a previously unappreciated role for WNT4 in regulating cellular respiration, in lipid and amino acid metabolism, and in metabolic remodeling of ILC and gynecologic cancers.

## RESULTS

### WNT4-BioID identifies distinct WNT4 intracellular localization to the mitochondria

Our prior work supports that intracellular WNT4 activity regulates mitochondrial function [6, 25], but WNT4 localization and signaling partners for intracellular signaling are unknown. To profile WNT4 trafficking, localization, and novel intracellular functions, we performed proximity-dependent biotinylation (BioID) followed by mass spectrometry (MS). We expressed WNT4 or WNT3A fused to C-terminal BirA biotin ligase in HT1080 wild-type (WT) and porcupine O-acetyltransferase (*PORCN*)-knockout (PKO) cells (Wnt-responsive fibrosarcoma cell line), and ILC cell line MDA MB 134VI (MM134) (**Supplemental Figure 1A-C**). Use of PKO knockout cells allowed us to validate that BioID with Wnt over-expression appropriately maps Wnt trafficking (discussed below), and we previously identified WNT4 intracellular signaling and trafficking in each of these models [6, 25]. Cells were biotin-treated for 24 hours, and biotinylated proteins - i.e. those within ∼10 nm of Wnt-BirA (**Supplemental Figure 1C**) - were extracted and profiled by MS as described in Methods. Raw data for BioID-MS is provided in **Supplemental File 1**.

We confirmed our BioID-MS approach could identify differential canonical WNT3A trafficking based on PORCN status (HT1080 vs HT1080-PKO; **Supplemental Figure 1D**). In canonical Wnt trafficking, PORCN is required for Wnt interaction with Wntless (WLS) in the endoplasmic reticulum, subsequent WLS-mediated Wnt trafficking through the Golgi, and Wnt secretion; PORCN ablation inhibits Wnt trafficking and secretion.

Comparing WNT3A-associated proteins in HT1080 vs HT1080-PKO (**Supplemental File 2**), n=46 putative PORCN-dependent WNT3A-associated proteins included WLS (enriched >20-fold in HT1080 vs HT1080-PKO, **Supplemental Figure 1E**), and were enriched for Golgi localization (GO Cellular Compartment: trans-Golgi network, p=0.0024). These enrichments confirm that BioID identified suppression of WNT3A trafficking to the Golgi in PORCN-knockout cells. Only n=8 WNT4-associated proteins were enriched in HT1080 vs HT1080-PKO, suggesting PORCN-knockout had little impact on WNT4 trafficking, consistent with our prior report of atypical (PORCN-independent) WNT4 trafficking [25].

We compared WNT4-associated (n=277) vs WNT3A-associated (n=184) proteins in HT1080 (**Figure 1A**). From the WNT4-specific proteins in HT1080 (n=172), we also identified “high-confidence” WNT4-associated proteins based on increased signal for WNT4-BioID, vs control and WNT3A-BioID, in at least 2 of 3 cell lines (n=72, **Figure 1A-B, Supplemental File 2**). From these subsets, we examined differential predicted localization using gene ontology and SubCell BarCode analysis [26] (**Supplemental File 2**). Consistent with canonical Wnt secretion, WNT3A-associated proteins were enriched for localization at the ER/Golgi (**Figure 1C**), and WNT3A predicted localization was in the secretory pathway (**Figure 1D**). In contrast, WNT4-associated proteins were enriched for mitochondrial proteins (**Figure 1C,Supplemental File 2**). Using SubCell barcode analysis, WNT4 predicted localization was in the cytosol or at the mitochondria (**Figure 1D**). High-confidence WNT4-associated proteins were also enriched for mitochondrial proteins and predicted WNT4 localization at the mitochondria (**Figure 1B-D**). High-confidence WNT4-associated proteins included mitochondrial membrane and matrix proteins (**Figure 1E**, red text) as well as proteins involved in mitochondrial biogenesis/dynamics (**Figure 1E**, pink text), supporting mitochondrial WNT4 localization.

**Figure 1.**
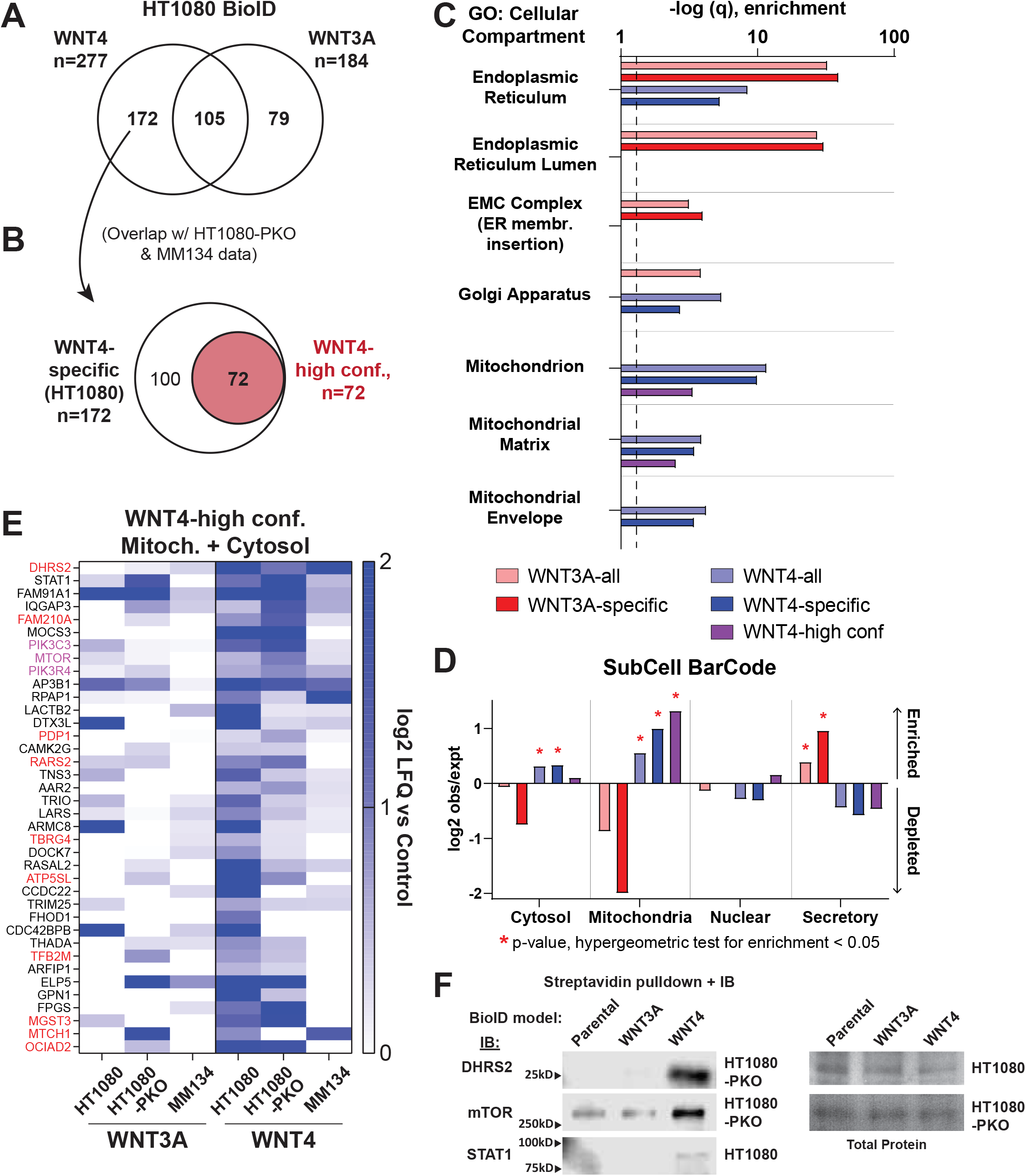
Proximity biotinylation (BioID) supports WNT4 localization to the mitochondria. (A) Proteins enriched in HT1080 Wnt-BirA versus parental HT1080 cells lacking BirA construct expression. (B) Overlap of proteins identified in HT1080 (panel A) versus HT1080-PKO and MM134 identifies n=72 “high-confidence” WNT4-associated proteins. (C) Gene ontology analysis for cellular compartment for WNT3A- vs WNT4-associated proteins. Dashed line = 1.3 (p = 0.05). (D) Network analysis of WNT3A- vs WNT4-associated proteins via Subcell Barcode. Enrichments against cell line HCC287 background shown; parallel results observed with other cell line background data, e.g. MCF7. (E) Proteins with predicted cytosolic or mitochondrial localization (subcell barcode) among “high-confidence” WNT4-associated proteins. Red = predicted mitochondrial localization, pink = mTOR complex in mitochondrial dynamics, biogenesis, and autophagy. (F) Biotin treatment and streptavidin pulldown was performed as for mass spectrometry studies, and candidate WNT4-associated proteins from (E) detected by immunoblotting. Total protein by Ponceau.

Among the most strongly enriched putative WNT4-associated proteins were mitochondrial reductase / dehydrogenase DHRS2, regulators of mitochondrial biogenesis, dynamics, and mitophagy (PIK3C3, PIK3R4, and mTOR), and STAT1, which can translocate to the mitochondria to regulate cellular metabolism [27–29]. For DHRS2, mTOR, and STAT1, we validated the putative association with WNT4 by streptavidin pulldown and immunoblotting in independent samples, i.e., pulldown of biotinylated DHRS2/mTOR/STAT1 was enriched in WNT4-BioID cells versus WNT3A-BioID or control cells (**Figure 1F**). These data support our BioID-MS findings and a novel role for WNT4 in regulating cellular metabolism and mitochondrial function.

### Integrated transcriptome and metabolome data support WNT4 as a mediator of ER metabolic signaling

As the mitochondria are essential for metabolic processes, we next examined how WNT4 controls metabolism in the context of dysregulated *WNT4* expression, using untargeted mass spectrometry-based metabolomics in ILC cell line MM134. MM134 are wild-type for the rs3820282 SNP (discussed below), but *WNT4* expression is aberrantly induced by ER in ILC cell lines - in the normal mammary gland and IDC cell lines, *WNT4* is directly PR controlled, independent of ER [3, 5]. **Figure 2A** summarizes the overall metabolomics study design. We targeted ER-WNT4 signaling directly using siRNA, and in parallel, compared inhibition of ER-WNT4 signaling in parental MM134 versus WNT4 over-expressing (W4OE) MM134 cells.

**Figure 2.**
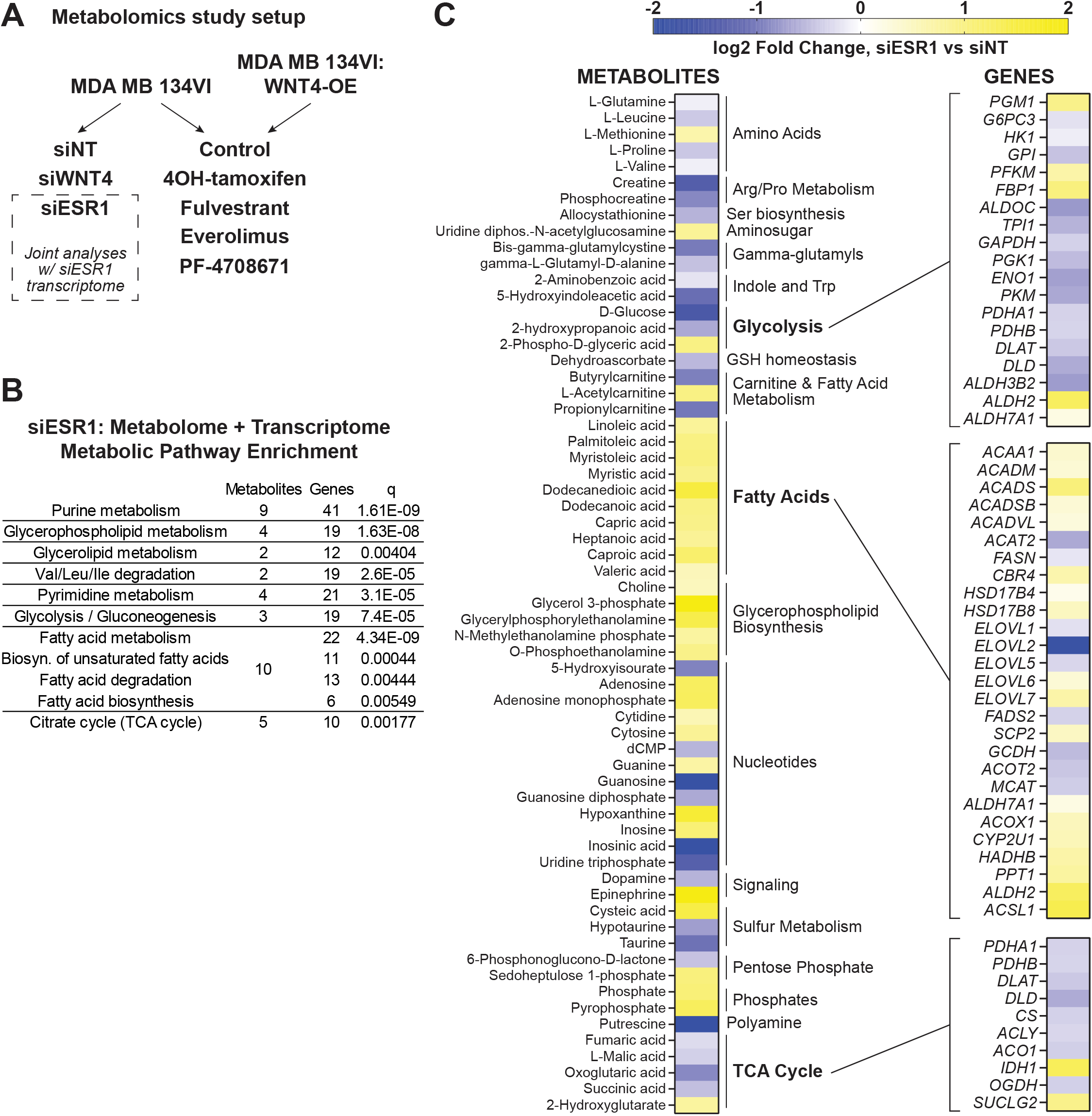
Estrogen receptor regulates glycolysis, oxidative phosphorylation, and fatty acid metabolism in ILC cells. (A) Overall metabolomics study design in MDA MB 134VI ILC cells; all samples in biological triplicate. (B) Joint analysis of trascriptome + metabolome data identifies dysregulated pathways after *ESR1* knockdown. Trascriptome data from GSE171364. (C) Metabolites levels altered by *ESR1* knockdown (left, n=63), with gene expression changes associated with major metabolic mechanisms (right).

To first examine the role for ER in regulation of metabolism in ILC cells, we integrated metabolomics data with transcriptomic data in MM134 (GSE171364 [30]), and performed joint pathway analysis of metabolites (n=63) and genes dysregulated by *ESR1* siRNA (n=5322) (**Figure 2B**; **Supplemental File 3**). Consistent with the central role for ER in cell proliferation, integrated pathway analysis showed enrichment in biosynthetic pathways, as well as glycolysis, the TCA cycle, and fatty acid metabolism (**Figure 2B**). Metabolite and gene expression changes after ER knockdown were consistent with decreased glycolysis and cellular respiration, and increased fatty acid metabolism (**Figure 2C**). Notably, while levels of most TCA Cycle metabolites and associated genes decreased upon ER knockdown, expression of *IDH1* and level of onco-metabolite 2-hydroxyglutarate increased after ER knockdown (no *IDH1/2* mutations have been reported in MM134 cells). Overall, these data are consistent with reports on the critical role for ER in regulating cellular metabolism in ER+ breast cancer cells and confirm metabolic remodeling upon suppression of ER.

We next compared effects of ER vs WNT4 knockdown (see Methods regarding siRNA validation). *WNT4* siRNA broadly dysregulated metabolite levels (n=77, **Figure 3A; Supplemental File 3**), consistent with our report of metabolic dysfunction upon WNT4 knockdown [6]. Metabolites dysregulated by WNT4 knockdown extensively mirrored ER knockdown (**Figure 3A-B**). Among n=46 metabolites dysregulated by both WNT4 and ER knockdown (e.g. fatty acids, 2-hydroxyglutarate, glutamine), siWNT4/siESR1 effects were strongly correlated (Spearman ρ = 0.8435, **Figure 3C**), supporting that WNT4 is a downstream mediator of ER-driven metabolic regulation. Pathway analysis confirmed that WNT4 and ER knockdown similarly impacted biosynthetic and metabolic pathways (**Figure 3D**), e.g. TCA Cycle, fatty acid biosynthesis, and glutamate metabolism. WNT4 knockdown impacted the levels of more metabolites in pathways including unsaturated fatty acid biosynthesis (pathway enrichment: siESR1, p=0.79; siWNT4, p=0.033) and glutamine/glutamate metabolism (siESR1, p=0.26; siWNT4, p=0.017) (**Supplemental File 3**). While the metabolic impact of ER and WNT4 knockdown are tightly correlated, these differences suggest that some metabolic activities of WNT4 signaling are distinct from global transcriptomic and metabolic impacts of ER.

**Figure 3.**
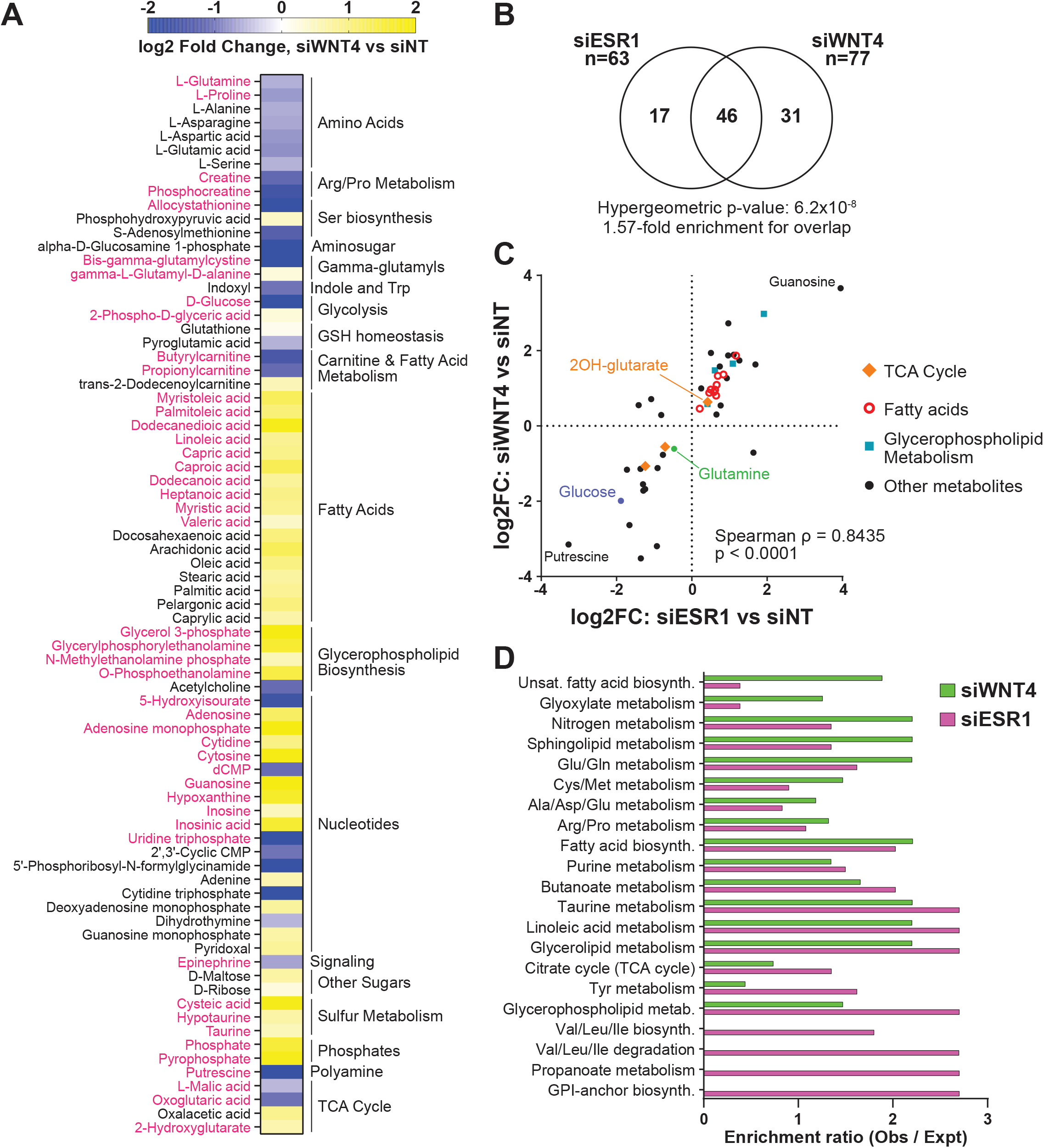
Metabolic effects of WNT4 knockdown mirror ER knockdown but have expanded impact on fatty acid and amino acid metabolic pathways. (A) Metabolite levels altered by *WNT4* knockdown (n=77); pink text indicates a shared affected metabolite with *ESR1* knockdown. (B) Metabolites dysregulated by *WNT4* v *ESR1* knockdown are strongly enriched for overlap. (C) *WNT4* v *ESR1* knockdown effects on metabolite levels are strongly directly correlated, suggesting an overall parallel effect on ILC cell metabolism. (D) Pathway analysis with *ESR1* metabolites (Figure 3C) and *WNT4* metabolites (Figure 4A); p < 0.4 in *ESR1* and/or *WNT4* dataset shown.

### WNT4 knockdown dysregulates cellular respiration but not glycolysis

We further examined the differential impact of ER vs WNT4 knockdown on cellular energetics using Seahorse metabolic flux analysis in MM134 cells (**Figure 4A**). Paralleling the observed effects on gene expression and metabolite levels in the TCA Cycle, *ESR1* or *WNT4* siRNA suppressed basal oxygen consumption rate (OCR; **Figure 4A-B**). However, with ER knockdown, cells retained full spare respiratory capacity, whereas WNT4 knockdown caused a decrease in spare respiratory capacity (**Figure 4B**); this observation supports that WNT4 knockdown causes mitochondrial dysfunction [6]. Unlike the effects of WNT4 knockdown on respiration, we noted siWNT4 had only a modest impact on extracellular acidification rate vs siESR1 (ECAR; **Figure 4C**). Accordingly, in our metabolomics data siESR1 reduced intracellular lactic acid levels while siWNT4 had no effect compared to control (**Figure 4D**), which we also observed measuring lactic acid secretion (**Figure 4E**). The differential effect on ECAR and lactic acid levels support that WNT4-mediated metabolic regulation is likely independent of glycolytic activity.

**Figure 4.**
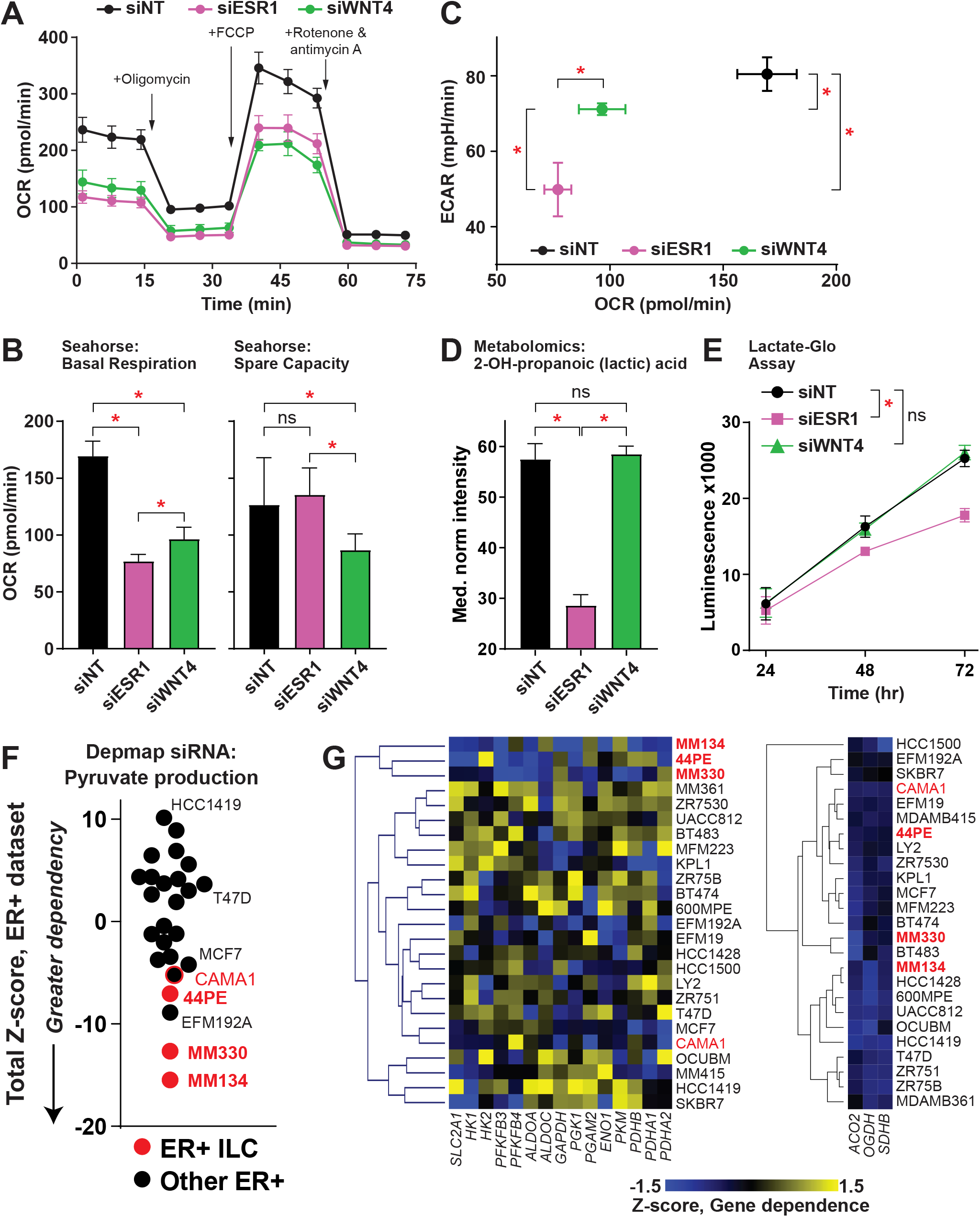
WNT4 knockdown impairs respiration but not glycolysis. (A) Seahorse MitoStress test in MM134. Points represent mean of 6 biological replicates ± SD. (B) Basal respiration is reduced by ER or WNT4 knockdown, but WNT4 knockdown impairs respiratory capacity. (C) WNT4 knockdown suppresses oxygen consumption (OCR), i.e. respiration (from B), but has a minimal effect on extracellular acidification (ECAR), i.e. glycolysis. (D) From metabolomics data, cellular lactic acid levels are reduced by ER knockdown, but not WNT4 knockdown. (B-D), comparisons by ANOVA with Dunnett’s correction. *, adj.p < 0.05. (E) Reduction in lactic acid production was also observed in lactate levels in conditioned medium. *, 2-way ANOVA siRNA effect p<0.05. (F-G) DEMETER2 scores for n=15 genes in pyruvate metabolism (KEGG M00001 & M00307) normalized as Z-scores for ER+ breast cancer cell lines. DEMETER2 siRNA used as ER+ ILC cell line data are not available in Depmap CRISPR-based screens. CAMA1 denoted separately as an “ILC-like” cell line. (F) Total sum of Z-scores for the geneset per cell line. (G) Heirarchal clustering for gene z-scores indicates ER+ ILC lines as an independent cluster. Pyruvate metabolism genes at left, representative TCA genes at right.

Since WNT4 knockdown suppresses ILC cell ATP levels and dysregulates oxygen consumption/respiration, but not glycolysis, we hypothesized that ILC cells are relatively more reliant on OXPHOS versus glycolysis for energy production. In cellular dependency screens (Depmap [31]), ILC cell lines are uniquely sensitive to suppression of genes involved in pyruvate metabolism. We examined 15 genes in glucose metabolism and pyruvate synthesis in 25 ER+ cell lines. ILC cell lines (n=3; MM134, 44PE, MM330) were among the most sensitive to knockdown of genes in this pathway (**Figure 4F**) and clustered independently from other ER+ lines (**Figure 4G**, left). ILC-like cell line CAMA1 [30, 32] was similarly sensitive to knockdown of pyruvate production genes (**Figure 4F**). This phenotype was specific for pyruvate production as TCA Cycle genes (e.g. *OGDH*) were broadly essential across ER+ breast cancer cell lines (**Figure 4G**, right). Taken together, WNT4 signaling is necessary for ILC cellular respiration; glycolysis is maintained with WNT4 knockdown but is insufficient to maintain ILC cell viability, as ILC cells are more dependent on cellular respiration via pyruvate production.

### WNT4 signaling is integrated in an ER-mTOR pathway and is critical for lipid metabolism

Our metabolomics study also compared small molecule inhibition of ER-WNT4 signaling on parental MM134 vs WNT4-overexpressing (W4OE) MM134 (**Figure 5A**). We hypothesized that WNT4 over-expression would rescue a subset of inhibitor-mediated metabolic effects and identify direct WNT4 signaling effects. Two-factor analysis (W4OE vs. Parental, controlled for drug effect) showed that WNT4-overexpression significantly altered the levels of 71 metabolites (**Figure 5B**), of which 38 overlapped with siWNT4. The effects of W4OE vs. siWNT4 were inverse for 58% of the 38 metabolites (n=22; **Figure 5C**), supporting direct WNT4 regulation of metabolites including glucose, glutamate, 2-hydroxyglutarate, and fatty acids (n=7) (**Supplemental File 4**).

**Figure 5.**
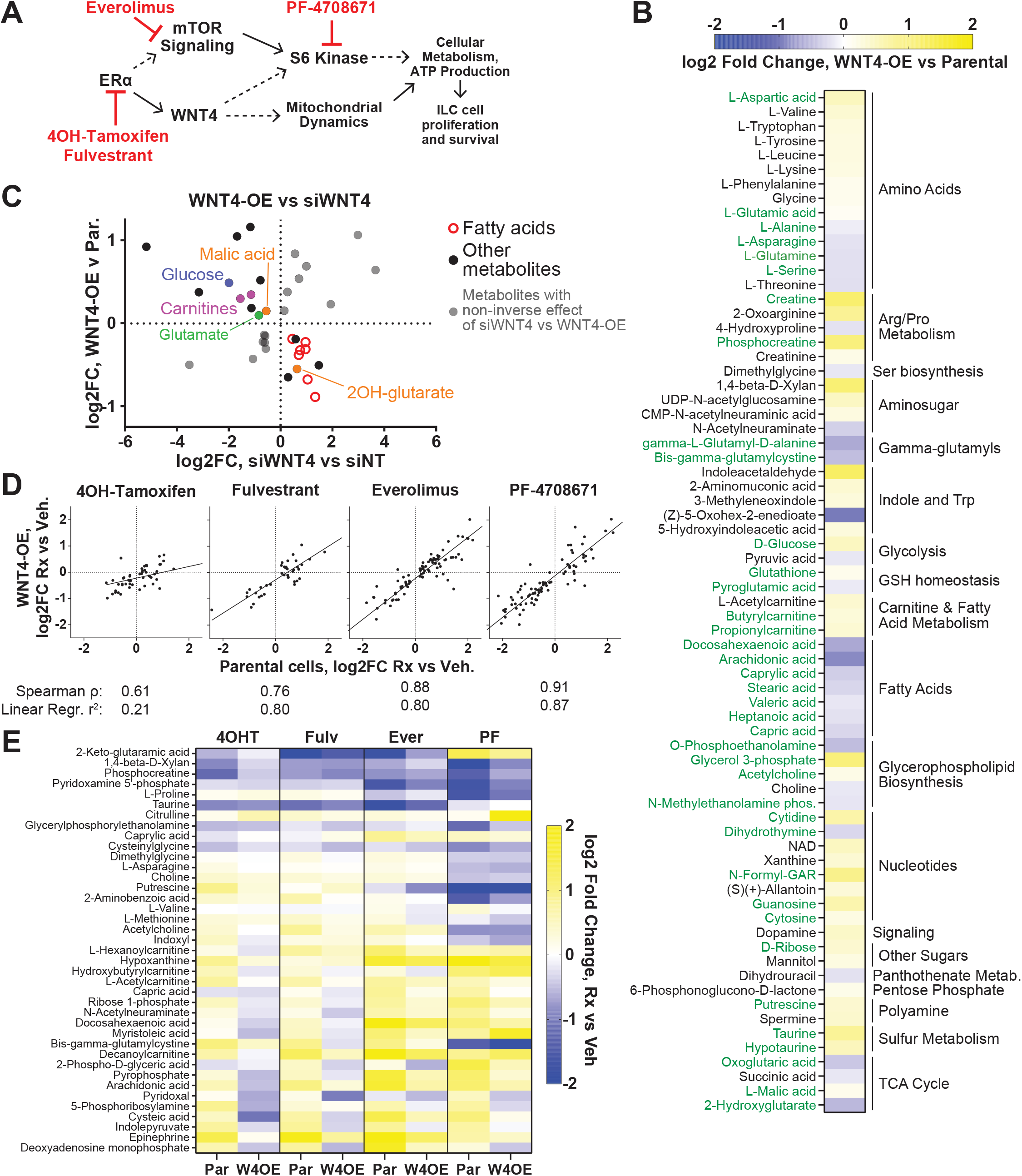
WNT4 over-expression rescues some metabolic effects of ER:WNT4 pathway inhibition. (A) Small molecule inhibitor targets in current model of ER:WNT4 signaling pathway. (B) Meta-analysis of parental MM134 cells versus WNT4-OE MM134 across drug treatment series (i.e. WNT4 effect controlled for inhibitor effect) identifies n=71 metabolite levels altered by WNT4 over-expression. Green = overlap with siWNT4 dysregulated metabolites. (C) Overall changes in metabolite levels caused by WNT4 knockdown vs WNT4 over-expression are inversely correlated, supporting regulation of associated metabolic pathways by WNT4. (D) Inhibitor effects in parental MM134 vs WNT4-OE MM134 are strongly correlated, but a subset of inhibitor effects are reversed by WNT4 over-expression. (E) Metabolites for which inhibitor effects is decreased by ≥30% for at least 2 inhibitors (n=39).

We further examined whether WNT4 over-expression rescued the metabolic effects of individual inhibitors. The effects of mTOR inhibitor everolimus and S6 Kinase inhibitor PF-4708671 on parental vs W4OE cells were strongly correlated (**Figure 5D**), suggesting that WNT4-overexpression has limited capacity to override inhibition of downstream signaling. Similarly, the effects of anti-estrogen fulvestrant were highly correlated in parental vs W4OE, as WNT4 may be insufficient to overcome complete ER inhibition in ER-dependent breast cancer cells. However, a larger subset of effects of 4OH-tamoxifen were reversed in W4OE cells, consistent with tamoxifen partial agonist activity in these cells [4]; incomplete ER inhibition by 4OH-tamoxifen (i.e. tamoxifen resistance) can be partially rescued by WNT4-overexpression. From these data, we identified n=38 metabolites for which effects of ≥2 inhibitors were reversed by ≥30% (**Figure 5E**), which were enriched for fatty esters (i.e. carnitines, n=4, p=0.032), supporting a key role for WNT4 in fatty acid/lipid metabolism.

Integrating WNT4-regulated metabolic pathways in siRNA, over-expression, and inhibitor studies identified 41 metabolites differentially regulated by activation vs suppression of WNT4 signaling (**Figure 6A, Supplemental File 5**). Enrichment analysis of these consensus WNT4-regulated metabolites identified the arginine and proline metabolic pathway, supporting a role for WNT4 signaling in glutamate metabolism. Fatty acid biosynthesis and metabolism were key pathways/metabolites regulated via WNT4 signaling (**Figure 6B**).

**Figure 6.**
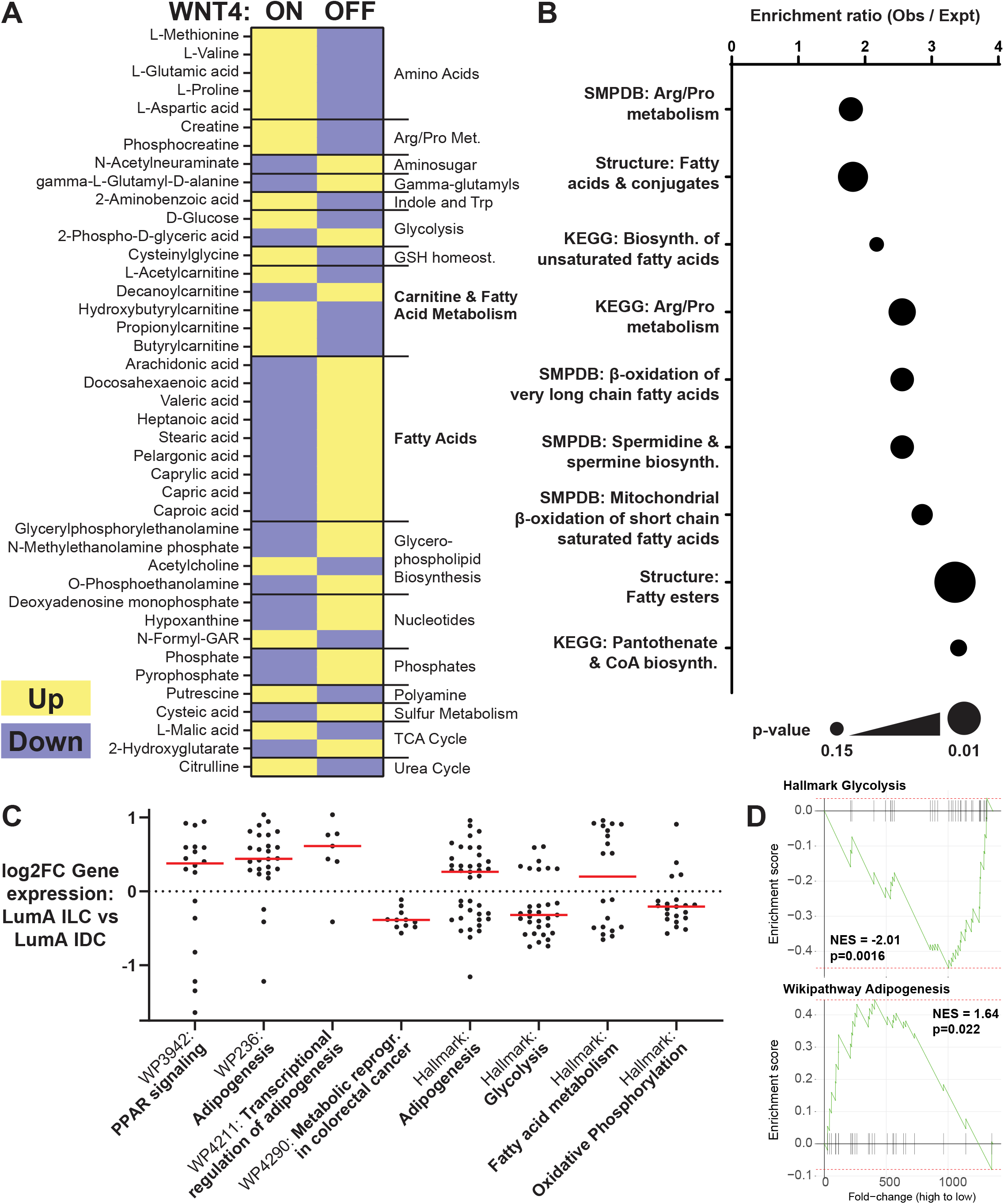
Consensus WNT4-regulated metabolites are enriched for fatty acid metabolism. (A) Consensus of n=41 metabolites rescued with WNT4-OE and metabolites inversely regulated by WNT4-OE vs WNT4 knockdown. WNT4 ON v OFF corresponds to differential regualtion by WNT4 over-expression vs knockdown, respectively. (B) Pathway analysis of consensus WNT4-regulated metabolites. (C) Gene set enrichment analysis of genes differentially expressed in Luminal A ILC vs Luminal A IDC (n=1360) includes metabolic pathways, with relative gene expression levels in ILC consistent with increased lipid metabolism and decreased glycolysis and OXPHOS. Points = individudal genes (fold-changes from TCGA, Ref 15); red bar = median fold change. (D) fGSEA of representative pathways in (C). NES = normalized enrichment score.

Our data on ER-WNT4 signaling highlights a putative role for lipid metabolism in the distinct metabolic phenotype of ILC. To examine this in tumor data, we performed gene set enrichment analyses from n=1360 genes differentially expressed in ER+ Luminal A ILC versus IDC [15]. Four of the top 8 over-represented pathways (Wikipathways) were metabolic, with an increase in genes associated with lipid metabolism in ILC (**Figure 6C, Supplemental File 6**). Metabolic pathways were also enriched in Hallmark gene signatures, with an increase in genes associated with Adipogenesis and Fatty acid metabolism, versus a decrease in genes associated with Glycolysis and Oxidative Phosphorylation, in ILC vs IDC (**Figure 6C**). Using functional gene set enrichment analyses, differential gene expression in ILC is consistent with decreased glycolysis and increased fatty acid metabolism (**Figure 6D, Supplemental File 6**). These data support that WNT4 signaling contributes to increased lipid metabolism as part of the distinct metabolic phenotype of ILC.

### Genetic polymorphism at the *WNT4* locus is associated with a distinct metabolic phenotype

We next examined rs3820282 genotype as a convergent mechanism of activating WNT4 signaling, paralleling ER regulation of WNT4 signaling in ILC, and based on the increased risk of gynecologic cancer associated with rs3820282 variant genotype [1], we expanded our studies into related models. We genotyped a panel of breast, ovarian, and endometrial cancer cell lines (n=58; **Supplemental File 7**). Among breast cancer cell lines, only 4 of 30 lines carried variant alleles (variant allele frequency ∼13%). In ovarian and endometrial cancer cell lines, 19 of 28 lines carried variant alleles (variant allele frequency ∼50%), likely reflecting both the increased diversity of patient ethnicity among these models and a distinct role for this SNP in gynecologic cancer. From these data we selected 4 wild-type and 4 variant genotype ovarian cancer cell lines to examine the impact of WNT4 knockdown, relative to our prior observations in ILC cells. *WNT4* siRNA reduced *WNT4* mRNA levels >70% in all cell lines (not shown), and strongly suppressed proliferation in rs3820282 variant cell lines but had modest or no effect in wild-type cell lines (**Figure 7A**). The differential effect of WNT4 knockdown suggests variant genotype cells are specifically dependent on WNT4 signaling. To examine activity of WNT4 signaling, we measured levels of MCL1 by immunoblot, which we showed increases in ILC cells upon WNT4 knockdown and correlates with mitochondrial dysfunction [6]. Consistent with the proliferation results, MCL1 levels specifically increased in variant genotype models after WNT4 knockdown (**Figure 7B**). Collectively these data support that in ovarian cancer cells, WNT4 signaling is specifically active in the context of rs3820282 variant genotype.

**Figure 7.**
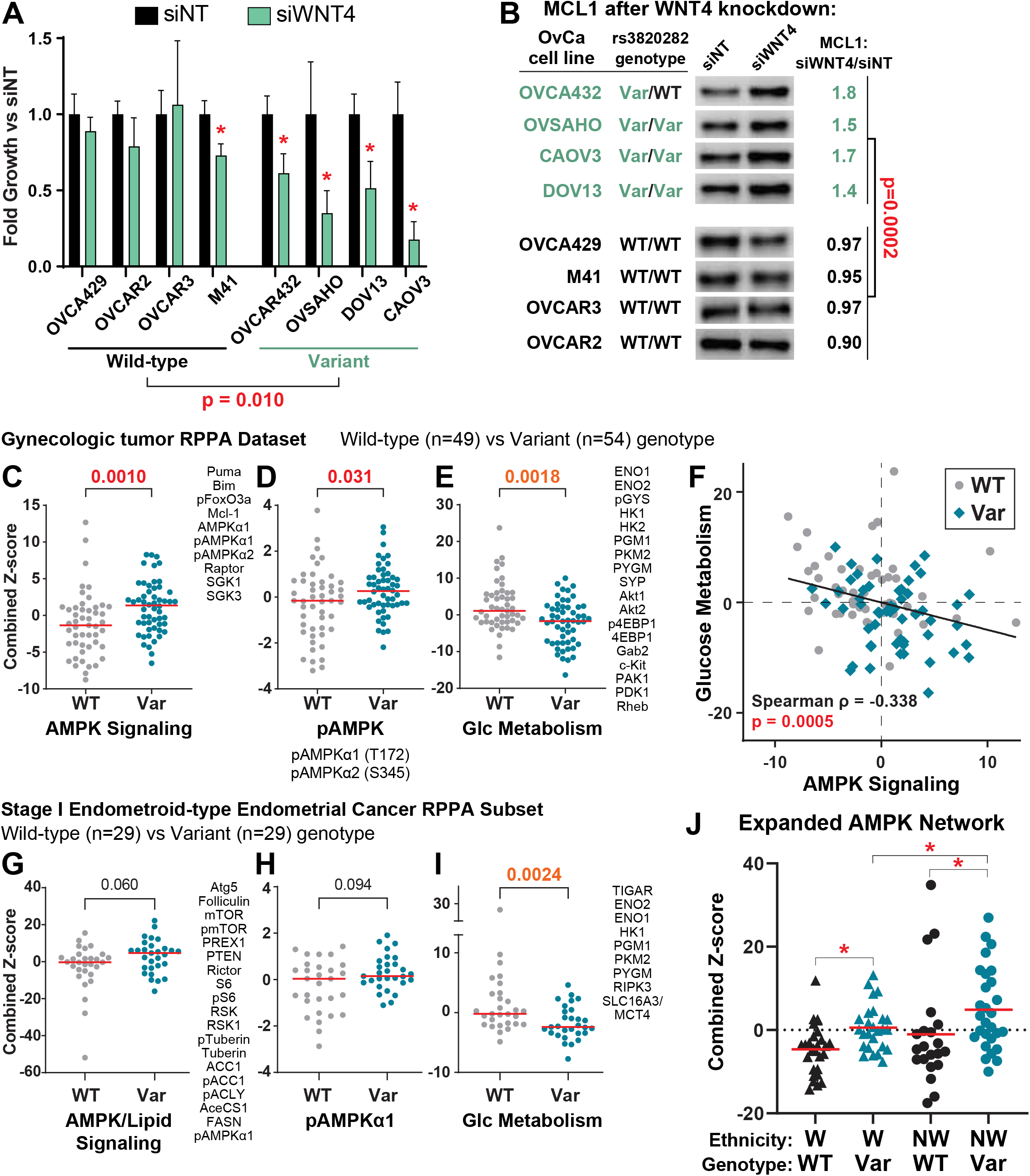
*WNT4* variant genotype is associated with active WNT4 signaling and metabolic remodeling in gynecologic tumors. **(A)** Proliferation assessed by dsDNA quantification 6 days post-transfection with siRNA (siNT = non-targeting control pool). Bars represent mean of 6 biological replicates ± SD; *, p<0.05, siWNT4 vs siNT, T-test with Welch’s correction. WT v Var model comparison by T-test of mean fold changes for siWNT4 vs siNT. **(B)** Lysates harvested 72hr post-transfection. MCL1 levels assessed by densitometry and normalized to total S6 loading control (not shown)[6]. Fold change in normalized MCL1 levels in Variant vs WT genotype cell lines compared by Student’s T-test. **(C-E)** Points represent total network score for the listed RPPA targets for individual tumors based on rs3820282 genotype. Red bar = median score; p-value from Mann-Whitney T-test. **(F)** Scatterplot of network scores in panels A and C; correlation from full dataset, i.e. WT and Var samples combined. **(G-I)** Network scores as in (A-C) from n=58 Stage I Endometriod-type endometrial tumors. **(J)** Points represent AMPK network score for individual tumors based on rs3820282 genotype and race/ethnicity groups. *, adj.p<0.1, ANOVA with Welch’s correction.

These observations led us to further explore the role of WNT4 in gynecologic tumor biology, also based on the key roles for WNT4 across reproductive tissues, the convergent mechanisms of *WNT4* dysregulation with rs3820282 vs ER in ILC, and parallels in ILC vs gynecologic cancer biology (see Discussion). We leveraged a biobank of snap frozen tissues with matched germline DNA samples via the U. Colorado Gynecologic Tumor & Fluid Bank (GTFB). With this resource, we were able to enrich for a diverse patient population to reflect increased variant allele frequency in Latinx and East Asian populations (see Discussion), and collect sufficient variant genotype tissues for analysis. We performed Taqman SNP genotyping for rs3820282 on genomic DNA samples from n=226 patients, and subsequently selected tissues from 103 patients (**Table 1**) for reverse phase protein array (RPPA) analyses. We enriched the RPPA cohort for germline variant genotypes (heterozygous and homozygous variant, 52%) and samples from non-White patients (47%; see Discussion). Of note, no significant differences in age or BMI were associated with rs3820282 genotype (**Table 1**).

**Table 1.**
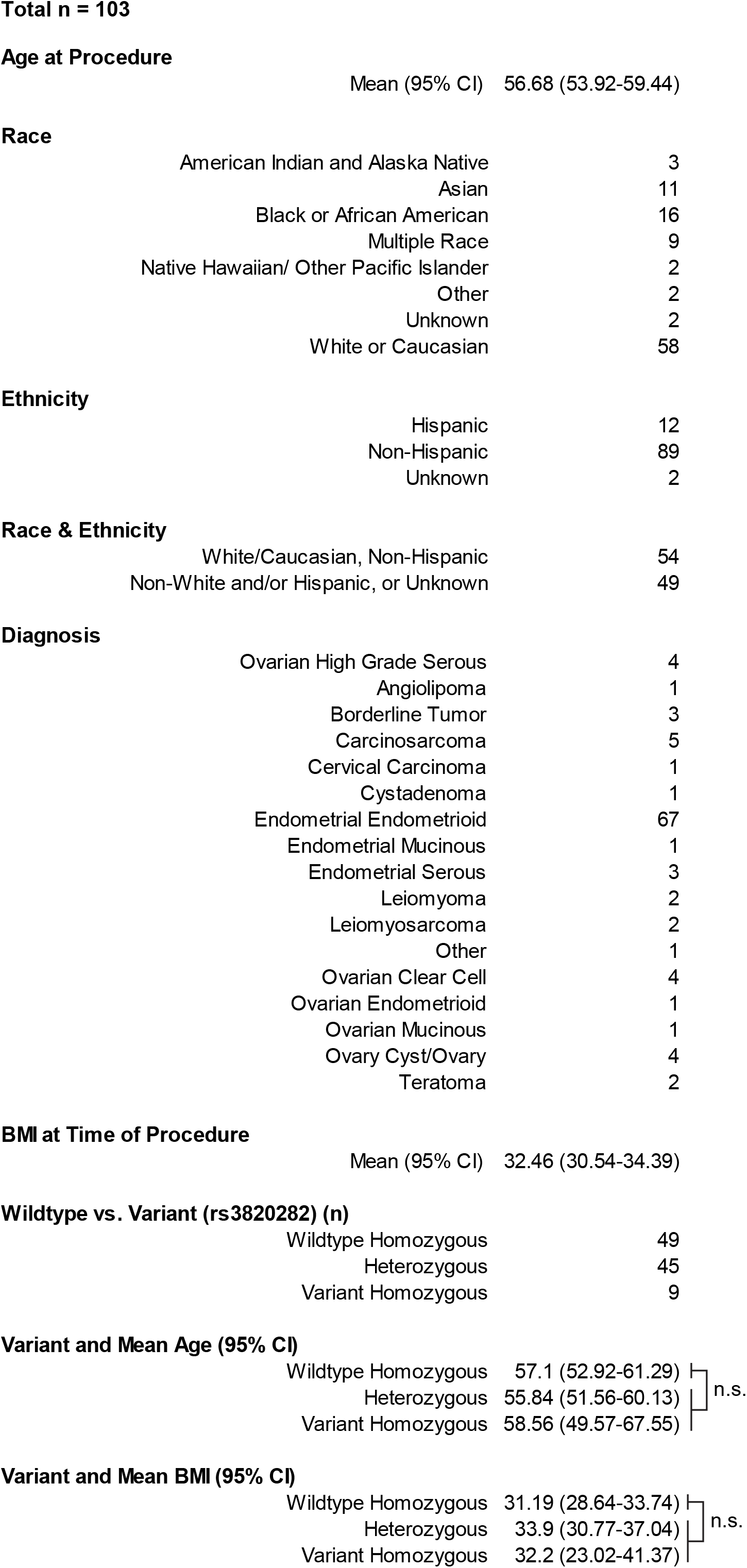
Gynecologic Tumor Cohort for RPPA Analyses.

**Race**
|  |  |
| --- | --- |
| American Indian and Alaska Native | 3 |
| Asian | 11 |
| Black or African American | 16 |
| Multiple Race | 9 |
| Native Hawaiian/ Other Pacific Islander | 2 |
| Other | 2 |
| Unknown | 2 |
| White or Caucasian | 58 |
| **Ethnicity** |  |
| Hispanic | 12 |
| Non-Hispanic | 89 |
| Unknown | 2 |
| **Race & Ethnicity** |  |
| White/Caucasian, Non-Hispanic | 54 |
| Non-White and/or Hispanic, or Unknown | 49 |
| **Diagnosis** |  |
| Ovarian High Grade Serous | 4 |
| Angiolipoma | 1 |
| Borderline Tumor | 3 |
| Carcinosarcoma | 5 |
| Cervical Carcinoma | 1 |
| Cystadenoma | 1 |
| Endometrial Endometrioid | 67 |
| Endometrial Mucinous | 1 |
| Endometrial Serous | 3 |
| Leiomyoma | 2 |
| Leiomyosarcoma | 2 |
| Other | 1 |
| Ovarian Clear Cell | 4 |
| Ovarian Endometrioid | 1 |
| Ovarian Mucinous | 1 |
| Ovary Cyst/Ovary | 4 |
| Teratoma | 2 |
| **Wildtype vs. Variant (rs3820282) (n)** |  |
| Wildtype Homozygous | 49 |
| Heterozygous | 45 |
| Variant Homozygous | 9 |

**Variant and Mean Age (95% CI)**
|  |  |  |
| --- | --- | --- |
| Wildtype Homozygous | 57.1 (52.92-61.29) | } n.s. |
| Heterozygous | 55.84 (51.56-60.13) |  |
| Variant Homozygous | 58.56 (49.57-67.55) |  |

**Variant and Mean BMI (95% CI)**
|  |  |  |
| --- | --- | --- |
| Wildtype Homozygous | 31.19 (28.64-33.74) | } n.s. |
| Heterozygous | 33.9 (30.77-37.04) |  |
| Variant Homozygous | 32.2 (23.02-41.37) |  |

RPPA data for 484 targets (**Supplemental File 8**) was examined for differentially expressed protein networks in tissues from wild-type vs variant genotype patients as described in **Supplemental File 8** and Methods. We identified the top 40% of RPPA targets based on differential levels in wild-type vs variant samples (See Methods; n=194). Using these targets for hierarchical clustering yielded two major clusters with similar distributions of samples based on tissue type, genotype, and race/ethnicity (**Supplemental Figure 2**). STRING network analyses [33] of the RPPA targets differentially increased in rs3820282 wild-type (n=100) vs variant samples (n=94) identified increased activity of 4 signaling networks in variant genotype tissues, and 5 networks in wild-type genotype tissues (**Supplemental File 8, Supplemental Figure 3**). Several networks converged on metabolic signaling, including activation of an AMPK signaling network in variant genotype tumors (**Figure 7A**). Activated AMPK (phospho-AMPKα1 and α2) alone was significantly increased in variant allele tumors (**Figure 7B**). Conversely, glucose metabolism networks were increased in wild-type tumors (**Figure 7C, Supplemental Figure 3**). From these networks, AMPK signaling was inversely correlated with glucose metabolism, with variant genotype tumors showing increased AMPK signaling with decreased glucose metabolism signaling (**Figure 7D**). These gynecologic tumor data parallel our observations in ILC and support that WNT4 underpins metabolic remodeling.

We also performed a parallel subset analysis of the largest tumor type in the cohort, stage I endometrioid-type endometrial tumors (n=58 tumors; n=190 differential RPPA targets in WT vs variant samples). As in the full cohort, variant genotype samples (n=29) showed increased AMPK signaling activation, albeit more modest, but the associated STRING network also included increased lipid metabolism proteins (**Figure 7E-F**). Wild-type tumors showed significantly increased glucose metabolism signaling as in the full cohort (**Figure 7G**).

With the unique strength of our protein array study in enrichment for tumor samples from non-White patients, we explored how the AMPK network linked to the rs3820282 variant was further linked to ethnic background. We confirmed that an expanded AMPK network (n=26 targets, **Supplemental File 8**) was elevated in variant genotype samples in both White (W) and non-White (NW) patient populations; however, AMPK network score was highest specifically in the non-White variant genotype tumors (**Figure 7H**). Individual network protein signals were overall higher in the non-White sample set (**Supplemental File 8**), e.g. pAMPKα2_S345 (variant allele non-White vs White: mean z-score difference = 0.63, ANOVA adj.p = 0.0195).

## DISCUSSION

WNT4 plays a unique role in organogenesis, development, and pathology in female reproductive tissues, but mechanisms of WNT4 dysregulation and downstream signaling are not broadly well understood, due in large part to myriad tissue-specific effects of WNT4 [1]. Our observations – integrating proteomic, metabolomic, and transcriptomic studies in cell lines, primary human tumor samples, and various public datasets – suggest that convergent mechanisms of WNT4 dysregulation, via estrogen receptor in ILC [5] and SNP rs3820282 in gynecologic cancers [22, 24], drive cancer metabolism. We show that WNT4 acts via a novel intracellular mechanism, localizing to the mitochondria instead of canonical secretory/paracrine signaling, ultimately leading to metabolic remodeling with a decrease in glucose metabolism but increase in lipid and/or amino acid metabolism.

WNT4 localization to the mitochondria as predicted by BioID links our prior discovery of atypical intracellular localization and function of WNT4 [25], with our findings on WNT4 regulation of mTOR signaling and mitochondrial dynamics [6]. In the latter study, we found WNT4 is integral for mTOR signaling via S6 Kinase; WNT4 was necessary for S6 Kinase and S6 phosphorylation, but not mTOR phosphorylation. In the current study, we identified mTOR as a WNT4-associated protein. BioID is limited in not distinguishing proximity vs direct interaction, but WNT4 may regulate mTOR interaction with or access to partners like S6 Kinase. Our observations indicate previously unreported mechanistic links between WNT4, DHRS2, and STAT1. Notably, both DHRS2 and STAT1 are implicated in mitochondrial function and metabolism [27–29, 34, 35]. Future studies will define the discrete localization of WNT4 at/within the mitochondria, trafficking mechanisms, and direct protein interactions. Defining various tissue-specific WNT4 activities as canonical, non-canonical, or atypical/intracellular will be key future directions.

In addition to common links to WNT4 in organogenesis and development, ILC and gynecologic cancers, including ovarian cancer, share intriguing parallels in their interactions with the microenvironment. Strikingly, ILC metastasize to unique sites relative to other breast cancer, in part mimicking the spread of gynecologic cancer; ILC can spread to the peritoneum, gastrointestinal tract, ovary, and endometrium [36–38]. Similarly, data from laboratory models, tumor studies, and clinical imaging studies support that ILC and gynecologic cancers have distinct metabolic phenotypes which preferentially utilize fuels beyond glucose. In ILC, limited FDG avidity in PET-CT imaging strongly suggests ILC tumors preferentially utilize fuels other than glucose, such as amino acids and/or fatty acids. Studies support that ILC cell lines are uniquely dependent on glutamate uptake and metabolism [19, 20]. Moreover, upregulation of *SREBP* and *FASN* are critical to anti-estrogen resistance in ILC (which also requires WNT4 [5]); endocrine resistant ILC cell lines were hypersensitive to SREBP knockdown or inhibition versus parental cells [18]. Similarly, ovarian cancer progression relies on fatty acid metabolism, and *FASN* upregulation in ovarian tumors correlates to shorter overall survival [39]. In tissue microarray analyses (ILC n=108, IDC n=584), nearly all ILC were positive for hormone-sensitive lipase (HSL/*LIPE*), which hydrolyzes triglycerides into free fatty acids, while this was uncommon in IDC (HSL+: 93% ILC v 15% IDC) [17]. Fatty acid transporter FABP4 was expressed in 32% of ILC vs <2% of IDC, supporting that ILC require greater fatty acid utilization than IDC. FABP4 expression in ovarian cancer also promotes disease progression and chemoresistance [39, 40]. Our analyses of WNT4 signaling support that WNT4 underpins fatty acid metabolism, yet limited data exist to define how this metabolic remodeling is initiated. Notably, recent work shows that the tumor microenvironment and metastatic niche can dramatically reprogram ER-driven cellular metabolism [41]. WNT4 signaling may integrate hormonal and/or microenvironmental signals to regulate metabolism in ILC and gynecologic cancers.

Germline WNT4 SNPs are linked to 10-25% increased risk for gynecologic pathologies including endometriosis, leiomyoma, and ovarian cancer (OvCa) [1]. Kuchenbaecker et al showed that *WNT4* SNPs (e.g. rs3820282) are associated with an odds ratio (OR) for overall OvCa of 1.11 (p=8×10^−7^), OR=1.09 for the endometrioid [EC] histotype (p=0.05), OR=1.12 for the serous [HGSC] histotype (p=6×10^−6^), and OR=1.24 for the clear cell [CCC] histotype (p=5×10^−4^) [42]. Critically, rs3820282 VAF varies across ethnic populations: 0-6% in Black populations; 12-20% in Caucasian populations; 10-40% in Latinx populations; 45-55% in East Asian populations [23]. Our data support that the SNP has implications beyond risk, in tumor biology including WNT4 signaling and metabolic remodeling. As such, *WNT4* genotype may drive disparities in OvCa risk, but also create opportunities for precision treatments targeting WNT4 signaling and/or metabolism based on genotype. Future study of WNT4 in OvCa biology (and other gynecologic cancers) are critical to link *WNT4* genotype to cancer risk, patient prognosis, therapy response, patient outcomes, and cancer health disparities.

WNT4 regulates breast and gynecologic cancer metabolism via a previously unappreciated intracellular signaling mechanism at the mitochondria, with WNT4 mediating metabolic remodeling favoring lipid metabolism. Understanding how WNT4 signaling is dysregulated, by estrogen and genetic polymorphism in ILC vs gynecologic cancers, offers new opportunities for defining tumor biology, precision therapeutics, and personalized cancer risk assessment.

## METHODS

### Cell culture

MDA MB 134VI (MM134; RRID:CVCL_0617), HT1080 (RRID:CVCL_0317), and HT1080-A11 (*PORCN*-knockout, -PKO) were maintained as described [4, 25]. WNT4 over-expressing models (W4OE) and WNT3A over-expressing models were previously described and cultured in the same conditions as parental cell lines [25]. Ovarian cancer cell lines, OVCA429 (RRID:CVCL_3936), OVCAR2 (RRID:CVCL_3941), OVCAR3 (RRID:CVCL_0465), 41M (RRID:CVCL_4993, known derivative of OAW28), OVCA432 (RRID:CVCL_3769), OVSAHO (RRID:CVCL_3114), DOV13 (RRID:CVCL_6774), CAOV3 (RRID:CVCL_0201), were maintained as described [39, 43]. Wnt-BirA fusion expressing lines (described below) were also cultured as the parental cells. All lines were incubated at 37°C in 5% CO_2_. Cell lines are authenticated annually via the University of Arizona Genetics Core cell line authentication service and confirmed mycoplasma negative every four months (most recently confirmed negative in March 2023). Authenticated cells were in continuous culture <6 months.

17β-Estradiol (E2; cat#2824), Z-4-hydroxytamoxifen (4OHT; cat#3412), and fulvestrant (fulv / ICI182,780; cat#1047) were obtained from Tocris Bioscience (Bio-Techne, Minneapolis, MN, USA) and dissolved in ethanol. Everolimus (evero; cat#11597) and PF-4708671 (PF; cat#15018) were obtained from Cayman Chemical (Ann Arbor, MI, USA) and dissolved in DMSO.

siRNAs were reverse transfected using RNAiMAX (ThermoFisher) according to the manufacturer’s instructions. All constructs are siGENOME SMARTpool siRNAs (Dharmacon/Horizon Discovery, Lafayette, CO, USA), nontargeting pool #2 (D-001206-14-05), human *WNT4* (M-008659-03-0005), human *ESR1* (M-003401-04-0010). Knockdown validation and siRNA efficacy in ILC cells for siWNT4 and siESR1 are previously described [5, 6]. Cell proliferation was assessed via dsDNA quantification by hypotonic lysis and Hoescht 33258 fluorescence as described [25].

### Proximity-dependent biotinylation (BioID)

WNT3A and WNT4 were sub-cloned by Gateway recombination from the open-source Wnt library (Addgene Kit #1000000022) to the MAC-Tag-C vector (Addgene #108077) [44], generating Wnt ORFs with C-terminal BirA fusion (Wnt-BirA). Wnt-BirA plasmids were packaged for lentiviral transduction, and stably transduced into MM134, HT1080, and HT1080-PKO as previously described [25]. For proximity biotinylation analyses, parental and Wnt-BirA cells were treated with 50μM biotin (MilliporeSigma B4501) for 24 hours. Cells were then lysed with lysis buffer with 1% Triton X-100 (RPPA buffer [6]), and biotinylated proteins were extracted from the lysates using Pierce Streptavidin Magnetic Beads (ThermoFisher 88816) per the manufacturer’s instructions.

### BioID - mass spectrometry analyses

For mass spectrometry analyses, cell lines (MM134, HT1080, HT1080-PKO; parental, WNT3A-BirA, and WNT4-BirA for each) were plated to 10cm plates in biological duplicate, and treated with 50μM biotin for 24 hours. Cells were lysed and extracted as above; sample/bead slurry was submitted for mass spectrometry analyses at the Central Analytical Mass Spectrometry Facility at the University of Colorado Boulder. Detailed description of protein extraction and mass spectrometry are provided in **Supplemental File 9**.

Label-free quantification (LFQ) data from MaxQuant were processed using LFQ-Analyst [45] to define differentially enriched proteins across Wnt-BioID studies. For comparisons, missing/zero values were imputed using Perseus-type or MinProb algorithms, with downstream hits required to be identified using both methods. Wnt-associated proteins were defined by increased LFQ vs control (parental cells without Wnt-BirA) with adjusted p <0.4. Proteins identified in >20% of studies in the CRAPome database [46] were excluded from further analyses. Gene ontology analyses were performed using Enrichr [47], DAVID [48], and SubCell BarCode ([26], NetWork multi-protein localization tool).

### Immunoblotting

Immunoblotting was performed as described [6, 25]. Blots were probed with Streptavidin-HRP (Cell Signaling #3999; RRID:AB_10830897) or antibodies used according to manufacturer’s recommendations: WNT4 (R&D Systems, MAB4751; RRID:AB_2215448); WNT3A (Novus Biologicals, MAB13242); anti-HA-HRP conjugate (Cell Signaling #2999; RRID:AB_1264166); DHRS2 (Sigma HPA053915; RRID:AB_2682307); mTOR (Cell Signaling #2983; RRID:AB_2105622); STAT1 (Sigma HPA000931; RRID:AB_1080100).; MCL1 (Cell Signaling #5453; RRID:AB_10694494). We note that recent lots of WNT4 MAB4751 show substantially increased non-specific background relative to our prior studies [6, 25], and can detect over-expressed WNT4 but no longer reliably detect endogenous WNT4.

### Metabolomics analyses

MM134 cells were transfected with siRNA for 72 hours prior to harvest; MM134 and MM134-W4OE were treated with vehicle (0.6% DMSO), fulvestrant (100nM), 4OHT (100nM), everolimus (100nM), or PF-4708671 (30μM) for 48 hours prior to harvest. At harvest, cells were trypsinized and counted, then washed three times in PBS, and frozen as dry cell pellets prior to processing at the University of Colorado Anschutz Medical Campus Cancer Center Mass Spectrometry Shared Resource Core Facility (RRID:SCR_021988). Metabolite extraction and mass spectrometry details are provided in **Supplemental File 9**. Metabolite data was analyzed using Metaboanalyst 5.0 [49]; parameters for specific analyses described in text and in ‘readme’ sheets in associated supplemental data files.

### Metabolic assays

XF96 (Agilent Technologies, 102417-100) extracellular flux assay kits were used to measure oxygen consumption (OCR) and glycolytic flux (ECAR) as described [50]. MM134 cells were reverse transfected with siRNA as above, and 24 hours later, cells were collected by trypsinization and re-plated to an XF96 microplate at 80k cells/well in 6 biological replicate wells per condition. 24 hours after re-plating (i.e. 48 hours after siRNA transfection), medium was replaced with Seahorse XF media (Agilent Technologies, 102353-100) and the plate was incubated at 37C for 30 min. After incubation, metabolic flux analyses were performed both at basal conditions and after injection of 5mg/ml oligomycin (Sigma-Aldrich, 871744), 2mM FCCP (Sigma-Aldrich, C2920), 5mM Antimycin A (Sigma-Aldrich, A8774), and 5mM rotenone (Sigma-Aldrich, R8875) (chemicals generously provided by the Jordan Laboratory, CU Anschutz). Outputs were normalized to total cell number.

Extracellular lactate was measured by Lactate-Glo assay (Promega, J5021). MM134 were reverse transfected with siRNA as above in 96-well plates (25k cells/well), and incubated for 6hr at 37C prior to time 0. At the indicated timepoints, 3μL of medium collected and diluted in PBS (1:250 dilution). At time course completion, samples were mixed 1:1 in assay sample buffer, and read per the manufacturer’s instructions.

### Gynecologic tumor cohort and reverse phase protein array

The University of Colorado has an Institutional Review Board approved protocol in place to collect tissue from gynecologic patients with both malignant and benign disease processes. All participants are counseled regarding the potential uses of their tissue and sign a consent form approved by the Colorado Multiple Institutional Review Board (COMIRB, Protocol #07-935). Patients are consented prior to surgery at a clinical visit. Primary gynecologic tissues are collected from patients undergoing surgical resection for a suspected gynecologic malignancy. Blood and tissues are evaluated in the Department of Pathology and then processed at the GTFB laboratory within 2 hrs of surgical resection. DNA is isolated from the blood using the DNeasy columns (Qiagen). DNA and tissues are snap frozen and stored at -80C. Specimens were selected from banked GTFB tissues based on patient ethnicity and tumor types.

Snap frozen tumors were sent to MD Anderson Functional Proteomics RPPA Core Facility (Houston, TX; RPPA Core Facility RRID:SCR_016649). Tumor protein lysates were arrayed on nitrocellulose-coated slides. Sample spots were probed with 484 unique antibodies and visualized by DAB calorimetric reaction to produce stained slides. Relative protein target levels were determined for each and designated as log2 intensities and subsequently median-centered. Level 4 data (i.e. normalized for loading and batch effects) was used for analyses. Differential target levels were assessed using Morpheus (https://software.broadinstitute.org/morpheus) Signal to Noise Analysis, further described in **Supplemental File 7**.

### Single nucleotide polymorphism genotyping

Cell line gDNA and tumor gDNA from the GTFB were genotyped for rs3820282 using TaqMan assay C 27521414_20 (ThermoFisher #4351379) and TaqMan genotyping master mix (ThermoFisher #4371353). Reactions were carried out on a ThermoFisher QuantStudio6 Real-Time PCR System, and genotype calls were made using QuantStudio analysis software.

## ACKNOWLEDGEMENTS

The authors wish to thank Dr. Julie Haines and the University of Colorado School of Medicine Metabolomics Core, and Dr. Chris Ebmeier and the Central Analytical Mass Spectrometry Facility at the University of Colorado Boulder, for their contributions to this project. We thank the University of Colorado Anschutz Medical Campus Cancer Center Pathology Shared Resource Biorepository Core Facility (RRID:SCR_021989) for support with tumor tissue banking. We thank Dr. Courtney Jones and the Craig Jordan Laboratory for technical assistance with Seahorse metabolic flux analyses. We acknowledge philanthropic contributions from Kay L. Dunton Endowed Memorial Professorship in Ovarian Cancer Research, the McClintock-Addlesperger Family, Karen M. Jennison, Don and Arlene Mohler Johnson Family, Michael Intagliata, Duane and Denise Suess, Mary Normandin, and Donald Engelstad.

