## Supplemental Figure 1-3 for "WNT4 regulates cellular metabolism via intracellular activity at the mitochondria in breast and gynecologic cancers"

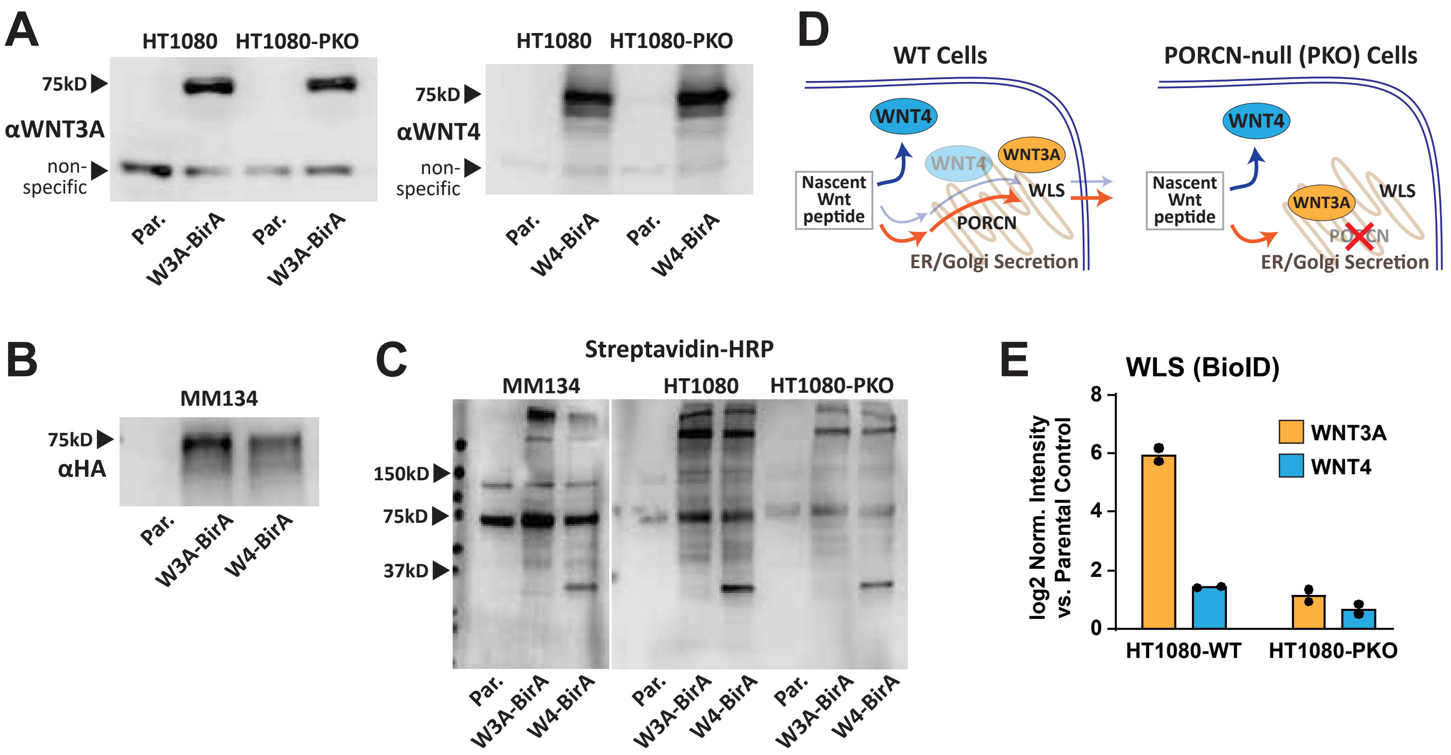


**Supplemental Figure 1. Wnt-BirA proximity biotinylation system differentiates PORCN-mediated Wnt trafficking.** (A-C) Immunoblotting of Parental cell lines (Par.) or cells stably expressing the indicated Wnt-BirA fusion protein. (A) Wnt-BirA fusions are predicted at ~75kD, and are detected by WNT3A or WNT4 antibodies at the expected size. Non-specific band shown as loading confirmation. (B) Wnt-BirA fusions can also be detected at the expected size using an internal HA-tag. (C) Parental and Wnt-BirA expressing cells were treated with 50μM biotin for 24hr, and whole cell lysates were analyzed for the presence of protein biotinylation using streptavidin-HRP conjugate. Wnt-BirA expressing show a broad range of protein biotinylation not observed in parental cells. (D) Model of WNT3A vs WNT4 trafficking in PORCN-wild type vs -knockout contexts. WNT4 trafficking to its intracellular localization is to be PORCN-independent, while WNT3A trafficking to the ER/Golgi and secretion is PORCN-dependent. (E) Mass spectrometry intensity for WLS in the indicated Wnt-BirA model versus parental cell control. Points represent biological duplicate samples.

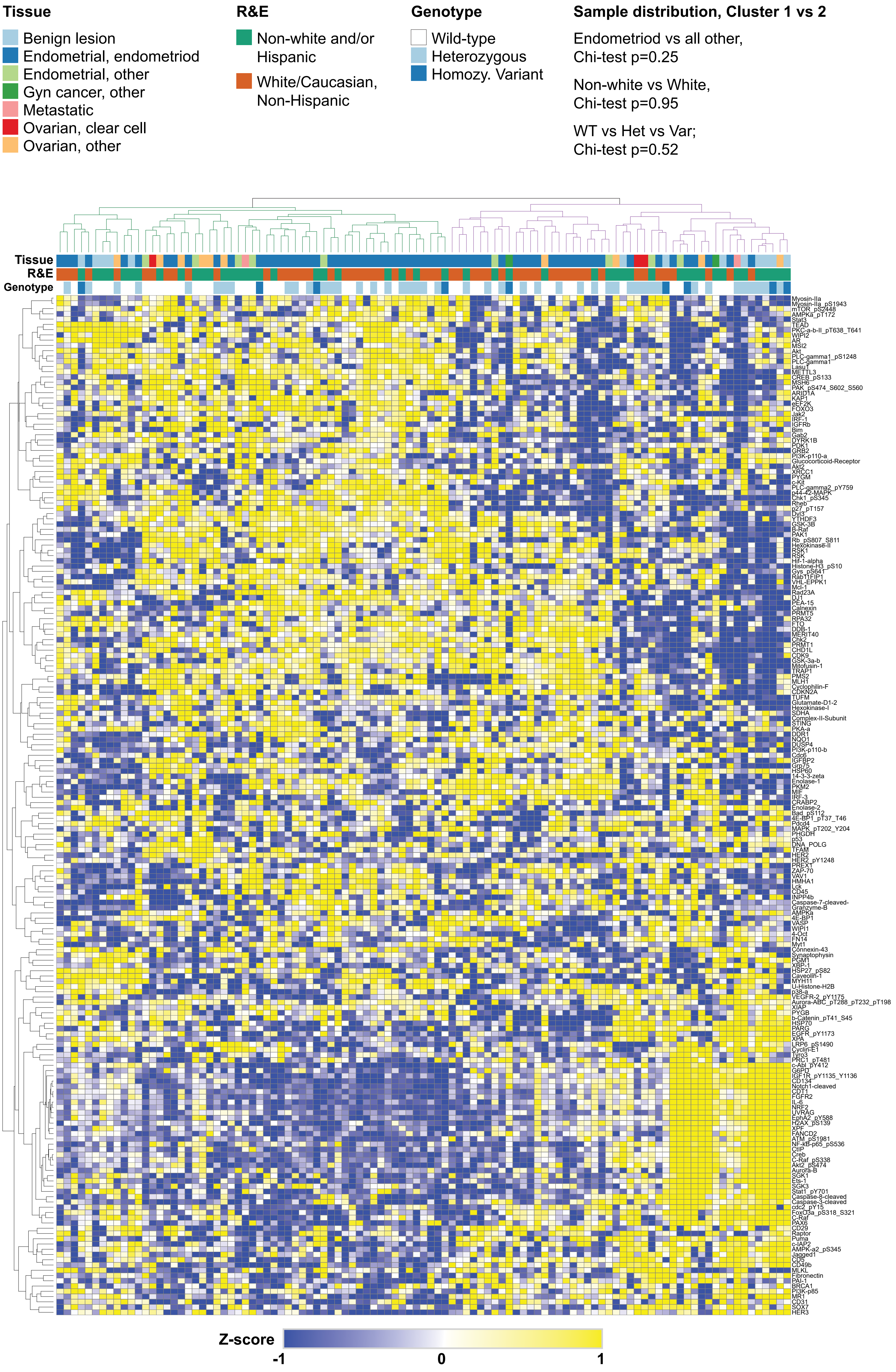


**Supplemental Figure 2. Differential protein array signals across gynecologic tissue cohort.** Top protein array targets based on differential levels in wild-type vs variant genotype tissues were subject to hierarchical clustering (Pearson) using Morpheus (see Methods). Two major sample clusters (Green vs Purple tree outlines at top) were not significantly different regarding distribution of tissue type, race & ethnicity, and rs3820282 genotype.

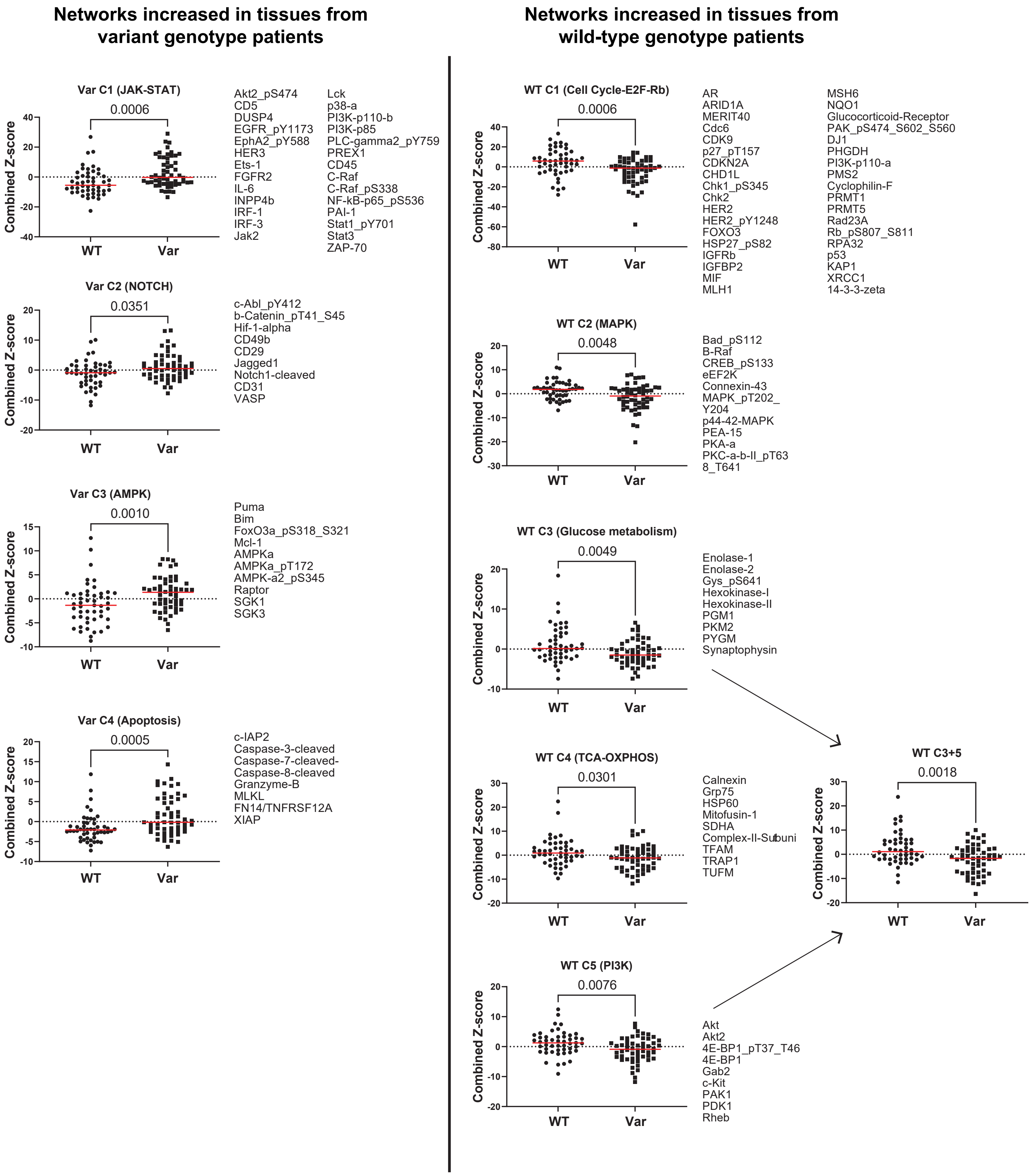


**Supplemental Figure 3. Protein signaling networks show differential activity by rs3820282 genotype.**
STRING networks defined by MCL clustering; networks shown with >6 protein array targets included (also see Figure 6). Points represent individual tissue samples and the sum of z-scores for the listed array targets. Red line = median score; comparisons by Mann-Whitney T-test.
