## Supplemental Files 1-9 for "WNT4 regulates cellular metabolism via intracellular activity at the mitochondria in breast and gynecologic cancers": Supplemental File 9 - Mass Spectrometry method supplement.docx

**Mass Spectrometry Methods**

**BioID – mass spectrometry analyses**

Samples were eluted from streptavidin beads, reduced, and alkylated with 2mM biotin in 5% (w/v) sodium dodecyl sulfate (SDS), 10 mM tris(2-carboxyethylphosphine) (TCEP), 40 mM 2-chloroacetamide, 50 mM Tris-HCl, pH 8.5 with boiling 10 minutes, then incubated shaking at 2000 rpm at 37°C for 30 minutes. Extracted proteins were digested using the SP3 method (1). Briefly, 200 µg carboxylate-functionalized speedbeads (Cytiva Life Sciences) were added followed by the addition of acetonitrile to 80% (v/v) inducing binding to the beads. The beads were washed twice with 80% (v/v) ethanol and twice with 100% acetonitrile. Proteins were digested in 50 mM Tris-HCl, pH 8.5, with 0.5 µg Lys-C/Trypsin (Promega) and incubated at 37˚C overnight. Tryptic peptides were desalted using an rpC18 column with 0.1% (v/v) TFA in water and acetonitrile. Cleaned-up peptide were then dried in a speedvac vacuum centrifuge and stored at -20°C until analysis.

Tryptic peptides were suspended in 3% (v/v) ACN, 0.1% (v/v) trifluoroacetic acid (TFA) and directly injected onto a reversed-phase C18 1.7 µm, 130 Å, 75 mm X 250 mm M-class column (Waters), using an Ultimate 3000 nanoUPLC (Thermos Scientific). Peptides were eluted at 300 nL/minute with a gradient from 2% to 20% ACN in 100 minutes then to 32% ACN in 20 minutes followed by 1 minute to 95% ACN and detected using a Q-Exactive HF-X mass spectrometer (Thermo Scientific). Precursor mass spectra (MS1) were acquired at a resolution of 120,000 from 380 to 1580 m/z with an automatic gain control (AGC) target of 3E6 and a maximum injection time of 45 milliseconds. Precursor peptide ion isolation width for MS2 fragment scans was 1.4 m/z, and the top 12 most intense ions were sequenced. All MS2 spectra were acquired at a resolution of 15,000 with higher energy collision dissociation (HCD) at 27% normalized collision energy. An AGC target of 1E5 and 100 milliseconds maximum injection time was used. Dynamic exclusion was set for 25 seconds with a mass tolerance of ±10 ppm. Rawfiles were searched against the Human database (UP000005640) using MaxQuant v.1.6.14.0. Cysteine carbamidomethylation was considered a fixed modification, while methionine oxidation and protein N-terminal acetylation were searched as variable modifications. All peptide/protein identifications were at a threshold of 1% false discovery rate (FDR).

**Metabolomics mass spectrometry**

Metabolites were extracted from frozen cell pellets at 3 million cells per mL by vigorous vortexing in ice cold 5:3:2 MeOH:MeCN:water (v/v/v) for 30 min at 4C. Supernatants were clarified by centrifugation (10 min @ 12kxg, 4C). The resulting extracts were analyzed (10uL per injection) by ultra-high-pressure liquid chromatography coupled to mass spectrometry (UHPLC-MS — Vanquish and Q Exactive, Thermo). Metabolites were resolved on a Kinetex C18 column (2.1 x 150 mm, 1.7 um) using a 5-minute gradient method as previously described (2). Following data acquisition, .raw files were converted to .mzXML using RawConverter then metabolites assigned and peaks integrated using Maven (Princeton University) in conjunction with the KEGG database and an in-house standard library. Quality control was assessed as using technical replicates run at beginning, end, and middle of each sequence as previously described (3).

1.	Hughes CS, Foehr S, Garfield DA, Furlong EE, Steinmetz LM, Krijgsveld J. Ultrasensitive proteome analysis using paramagnetic bead technology. Mol Syst Biol. 2014;10:757.

2.	Nemkov T, Reisz JA, Gehrke S, Hansen KC, D’Alessandro A. High-Throughput Metabolomics: Isocratic and Gradient Mass Spectrometry-Based Methods. Methods Mol Biol. 2019;1978:13–26.

3.	Nemkov T, Hansen KC, D’Alessandro A. A three-minute method for high-throughput quantitative metabolomics and quantitative tracing experiments of central carbon and nitrogen pathways. Rapid Commun Mass Spectrom. 2017;31:663–73.
